# Frequent genetic exchanges revealed by a pan-mitogenome graph of a fungal plant pathogen

**DOI:** 10.1101/2024.06.19.599757

**Authors:** Anouk C. van Westerhoven, Jelmer Dijkstra, Jose L. Aznar Palop, Kyran Wissink, Jasper Bell, Gert H. J. Kema, Michael F. Seidl

## Abstract

Mitochondria are present in almost all eukaryotic lineages. The mitochondrial genomes (mitogenomes) evolve separately from nuclear genomes, and they can therefore provide relevant insights into the evolution of their host species. *Fusarium oxysporum* is a major fungal plant pathogen that is assumed to reproduce clonally. However, horizontal chromosome transfer between strains can occur through heterokaryon formation, and recently signs of sexual recombination have been observed. Similarly, signs of recombination in *F. oxysporum* mitogenomes challenged the prevailing assumption of clonal reproduction in this species. Here, we construct, to our knowledge, the first fungal pan-mitogenome graph of nearly 500 *F. oxysporum* mitogenome assemblies to uncover the variation and evolution. In general, the gene order of fungal mitogenomes is not well conserved, yet the mitogenome of *F. oxysporum* and related species are highly co-linear. We observed two strikingly contrasting regions in the *Fusarium oxysporum* pan-mitogenome, comprising a highly conserved core mitogenome and a long variable region (6-16 kb in size), of which we identified three distinct types. The pan-mitogenome graph reveals that only five intron insertions occurred in the core mitogenome and that the long variable regions drive the difference between mitogenomes. Moreover, we observed that their evolution is neither concurrent with the core mitogenome nor with the nuclear genome. Our large-scale analysis of long variable regions uncovers frequent recombination between mitogenomes, even between strains that belong to different taxonomic clades. This challenges the common assumption of incompatibility between genetically diverse *F. oxysporum* strains and provides new insights into the evolution of this fungal species.

**Importance statement:** Insights into plant pathogen evolution is essential for the understanding and management of disease. *Fusarium oxysporum* is a major fungal pathogen that can infect many economically important crops. Pathogenicity can be transferred between strains by the horizontal transfer of pathogenicity chromosomes. The fungus has been thought to evolve clonally, yet recent evidence suggests active sexual recombination between related isolates, which could at least partially explain the horizontal transfer of pathogenicity chromosomes. By constructing a pan-genome graph of nearly 500 mitochondrial genomes, we describe the genetic variation of mitochondria in unprecedented detail and demonstrate frequent mitochondrial recombination. Importantly, recombination can occur between genetically diverse isolates from distinct taxonomic clades and thus can shed light on genetic exchange between fungal strains.

## Introduction

Mitochondria are the energy producing organelles of eukaryotic cells (Spinelli and Haigis 2018). Part of their genetic information is encoded apart from the nuclear DNA in a separate mitochondrial genome. This mitochondrial genome evolves at a different rate than the nuclear genome and behaves differently during cell division and reproduction (Barr et al. 2005; Burger et al. 2003; Smith and Keeling 2015). This results in different evolutionary dynamics that make the mitochondrial genome a useful phylogenetic marker which can help to elucidate evolutionary histories separate from the nuclear genome (Christinaki et al. 2022; Ingman et al. 2000).

Although the functions of mitogenomes in eukaryotes are largely conserved, their evolutionary dynamics vary between different eukaryotic lineages, with different mechanisms of inheritance, recombination rates, and mutation rates (Smith and Keeling 2015). In most species, mitochondria are uniparentally inherited, but the mechanisms resulting in uniparental inheritance vary between species and cases of biparental inheritance are also observed (Barr et al. 2005). In most bilateria, the mitochondrial structure is highly conserved and shows little signs of recombination (Ballard and Rand 2005; Ingman et al. 2000). In plants and fungi on the other hand, mitogenomes undergo extensive structural variation and show signs of recombination, horizontal transfer, and intron presence-absence variation (Fritsch et al. 2014; Megarioti and Kouvelis 2020; Smith and Keeling 2015). In fungal mitogenomes, recombination results in highly variable gene order, even within major fungal lineages (Aguileta et al. 2014). In addition to recombination, intron movement drives the structural evolution of fungal mitochondria. Mitochondrial introns are self-splicing mobile elements that can spread throughout and between genomes (Lang et al. 2007; Megarioti and Kouvelis 2020). In many fungal genomes, the expansion of introns is found to cause mitogenome size variations, and introns are often reported to be horizontally transferred (Repar and Warnecke 2017; Wu et al. 2015). These dynamics can provide useful evolutionary insights, for example intron activity can serve as a phylogenetic marker (Wang et al. 2021) and the recombination between mitochondria has revealed signs of introgression in *Saccharomyces* (Christinaki et al. 2022; Leducq et al. 2017; Wu et al. 2015). To further uncover the surprisingly different mechanisms of mitogenome evolution, in depth mitogenome analysis is needed for various species.

Research on mitochondrial evolution has mainly relied on comparing mitochondrial sequences to a reference genome (De Chiara et al. 2020) and as a result highly variable regions in mitochondrial genomes are difficult to analyse. Steps have been taken towards reference free mitogenome analysis using a pan-genomics approach (Brankovics et al. 2018; Wang et al. 2022a), however, large-scale pan-mitogenome analysis have, to our knowledge, not yet been conducted. Recent advances in pangenome variation graphs now allow for reference-free analysis of a large sets of genomes (Eizenga et al. 2020; Sherman and Salzberg 2020), providing a more complete view of gene presence / absence and structural variants (Hickey et al. 2020; Wang et al. 2022b). A pangenome variation graph stores sequences in many different nodes and each genome is represented as a path through connected nodes in the graph. Such a pan-mitogenome provides an excellent tool to analyse mitogenome evolution and enables the reconstruction of structural variants, intron homing, and possibly unique sequences obtained through horizontal transfer events.

The *Fusarium oxysporum* species complex contains several plant pathogens that infect many economically important crops (Edel-Hermann and Lecomte 2019). *F. oxysporum* is well known to exchange accessory chromosomes (Henry et al. 2021; Li et al. 2020; van Westerhoven et al. 2024) and importantly the horizontal transfer of specific accessory chromosomes can transfer pathogenicity between strains (Ma et al. 2010). Between which species this transfer can occur and how often this transfer occurs in nature remains unknown. *F. oxysporum* is considered an asexual fungus, however traces of meiotic recombination in a *F. oxysporum* population have recently been observed, indicating that *F. oxysporum* might in fact undergo sexual recombination (Fayyaz et al. 2023; McTaggart et al. 2021), which would be highly relevant to understand the evolution of this fungal species. The mitochondrial genome of *F. oxysporum* carries a conserved region encoding 14 core mitochondrial genes and a long variable region (LVR) (Al-Reedy et al. 2012; Brankovics et al. 2017). Interestingly, the distribution of this LVR is incongruent with the nuclear phylogeny, and thus mitogenomes seem to undergo recombination (Brankovics et al. 2017). This mitochondrial recombination supports the presence of a (para-)sexual cycle that involves the fusion of cells and at least the exchange of mitochondria. To further elucidate the evolutionary history of *F. oxysporum*, we here aim to analyse a large panel of mitogenomes and shed light on the mitochondrial recombination between strains. To this end, we constructed a pangenome variation graph of nearly 500 *F. oxysporum* mitogenomes to demonstrate frequent and ongoing genetic exchange between distantly related mitochondria, even between strains that belong to distinct taxonomic clades within the *F. oxysporum* species complex. These findings have direct implications for our understanding of the exchange and spread of genetic material, such as pathogenicity chromosomes, between strains of this economically and ecologically important group of fungi.

## Results

### Fusarium oxysporum mitogenomes have a highly conserved gene order

To understand the diversity of fungal mitogenomes, we first compared the mitogenomes of 706 taxonomically diverse fungal isolates (Fonseca et al., 2021, Table S1), including five *F. oxysporum* mitogenomes. As expected, the gene order is variable between fungal phyla, but is more similar to more closely related fungal genera (Fig. 1a), which corroborates previous findings (Aguileta et al. 2014). Unanticipatedly, mitochondrial gene order is conserved within the class of Sordariomycetes, to which also *F. oxysporum* belongs (Fig. 1a). Next to gene order variation, we also observed a large variation in mitogenome sizes ranging from only 12 kb in *Rozella allomycis* to up to 272 kb in *Ophiocordyceps camponoti-floridani* (Fig. 1b). Importantly, mitogenome size strongly correlates with intron abundance (Fig. 1c, Pearson r =0.76, p <0.05), indicating that recombination and intron expansions drive the variation of fungal mitogenomes.

**Figure 1.**
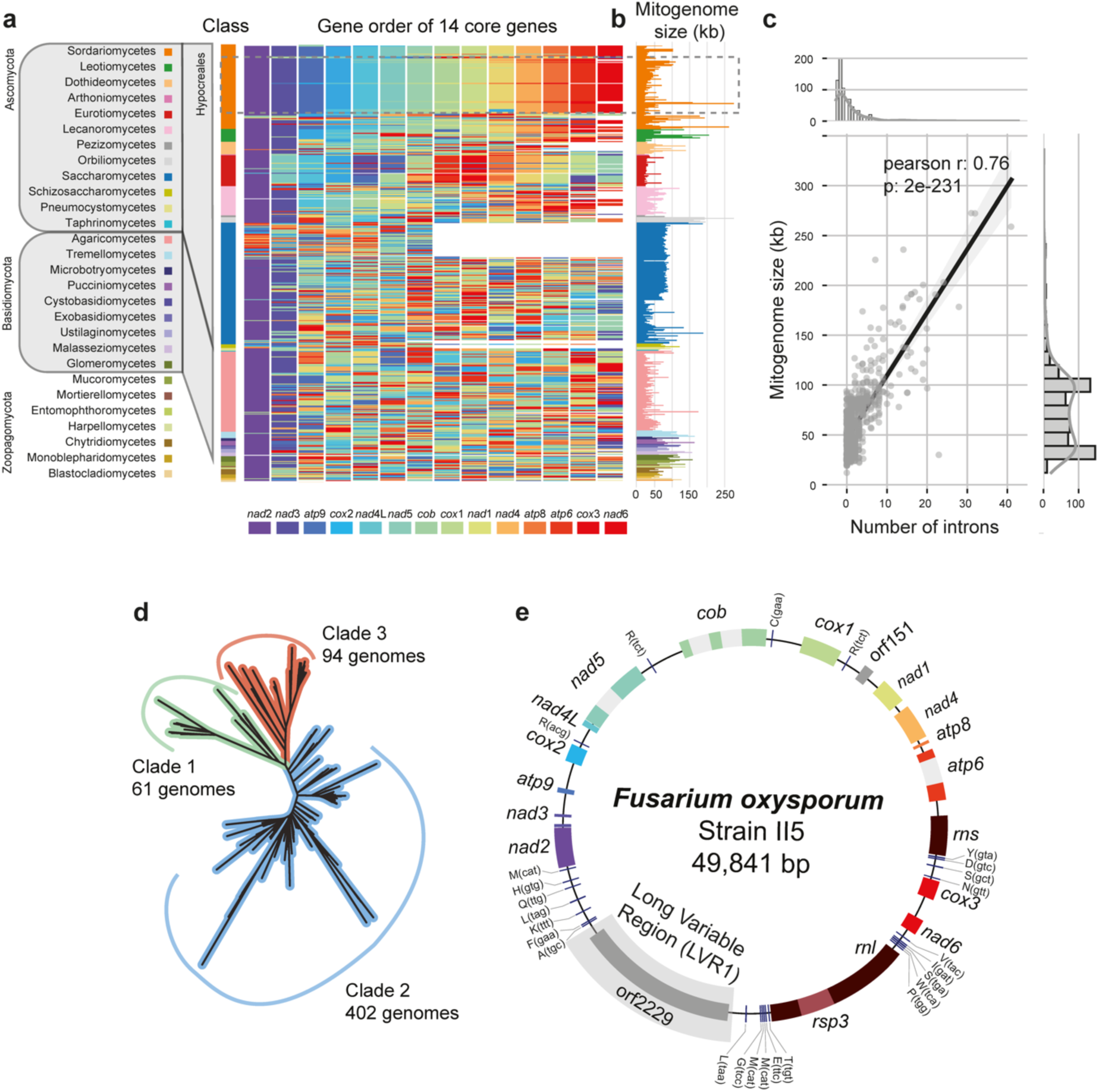
– Mitochondrial genome sizes and synteny are highly variable across fungi. (**a**) Gene order of 706 fungal mitochondria is highly variable, even between genetically closely related isolates. In contrast, mitochondria from fungi belonging to the Sordariomycetes are remarkably similar. *Fusarium oxysporum* belongs to the Hypocreales within the class of Sordariomycetes and have a highly conserved mitochondrial gene order. The 14 core genes of *Fusarium* mitochondria (Al-Reedy et al. 2012) are used to visualize the gene order. The mitochondrial genomes are ordered based on the fungal species phylogeny, and the coloured box indicates the identity of the gene and the respective position within the mitogenome; white boxes indicate missing genes. **(b)** Mitochondrial genomes are highly variable in size, even between closely related species **(c)** Mitochondrial genome size variation is largely driven by the number of introns. **(d)** Phylogenetic relationship of the publicly available *F. oxysporum* isolates used in this study. The phylogenetic tree was estimated based on 4213 conserved BUSCO genes using FastTree. **(e)** The mitochondrial genome assembly of *F. oxysporum* strain II5 (also known as *F. odoratissimum* strain II5 (Maryani et al. 2019; van Westerhoven et al. 2024) is shown as an example for mitogenomes in the *F. oxysporum* species complex. This mitogenomes contains all 14 core genes (coloured boxes), ribosomal proteins (dark red), four introns (grey boxes), tRNAs (blue bars) and a long variable region (LVR1).

To study the diversity and evolution of mitochondrial genomes in *F. oxysporum*, we obtained short-read sequencing data of 883 isolates (Table S2). We subsequently removed all isolates that do not belong to the *F. oxysporum* species complex and those where nuclear genome assemblies yielded low BUSCO completeness (Table S2). The remaining 618 *F. oxysporum* isolates are derived from 63 different projects (Table S2) and span the three previously described *F. oxysporum* taxonomic clades (O’Donnell et al. 1998), 61 genomes from clade 1, 402 from clade 2, and 94 from clade 3 (Fig. 1c), and thus represent a wide variety of *F. oxysporum* lineages.

We assembled the mitochondrial DNA using grabB (Brankovics et al. 2016), which resulted in complete and continuous circular mitogenome assemblies for 472 isolates (Fig. 2a, Table S2). The mitochondrial genome assemblies range between 37 and 55 kb in size (Fig. 2b, average 46 kb, Table S2) and have an average GC-content of 31.79%, which is in line with previous assemblies (Al-Reedy et al. 2012; Brankovics et al. 2017). For example, the mitochondrial genome of strain II5 is 49 kb and encodes the 14 *F. oxysporum* core genes, the small subunit RNA gene, the large subunit RNA gene, ribosomal protein S3 and six additional open reading frames (ORFs) (Fig. 1d). Four of the ORFs are part of an intronic homing endonuclease domain (Megarioti and Kouvelis 2020), but the role of the other two ORFs (ORF151, and ORF2229) is currently unknown. In total, between two and 17 additional ORFs were identified per assembled genome and the ORFs can be clustered into 77 families, indicating that the pan-mitogenome contains 77 ORFs in addition to the conserved core genes

**Figure 2.**
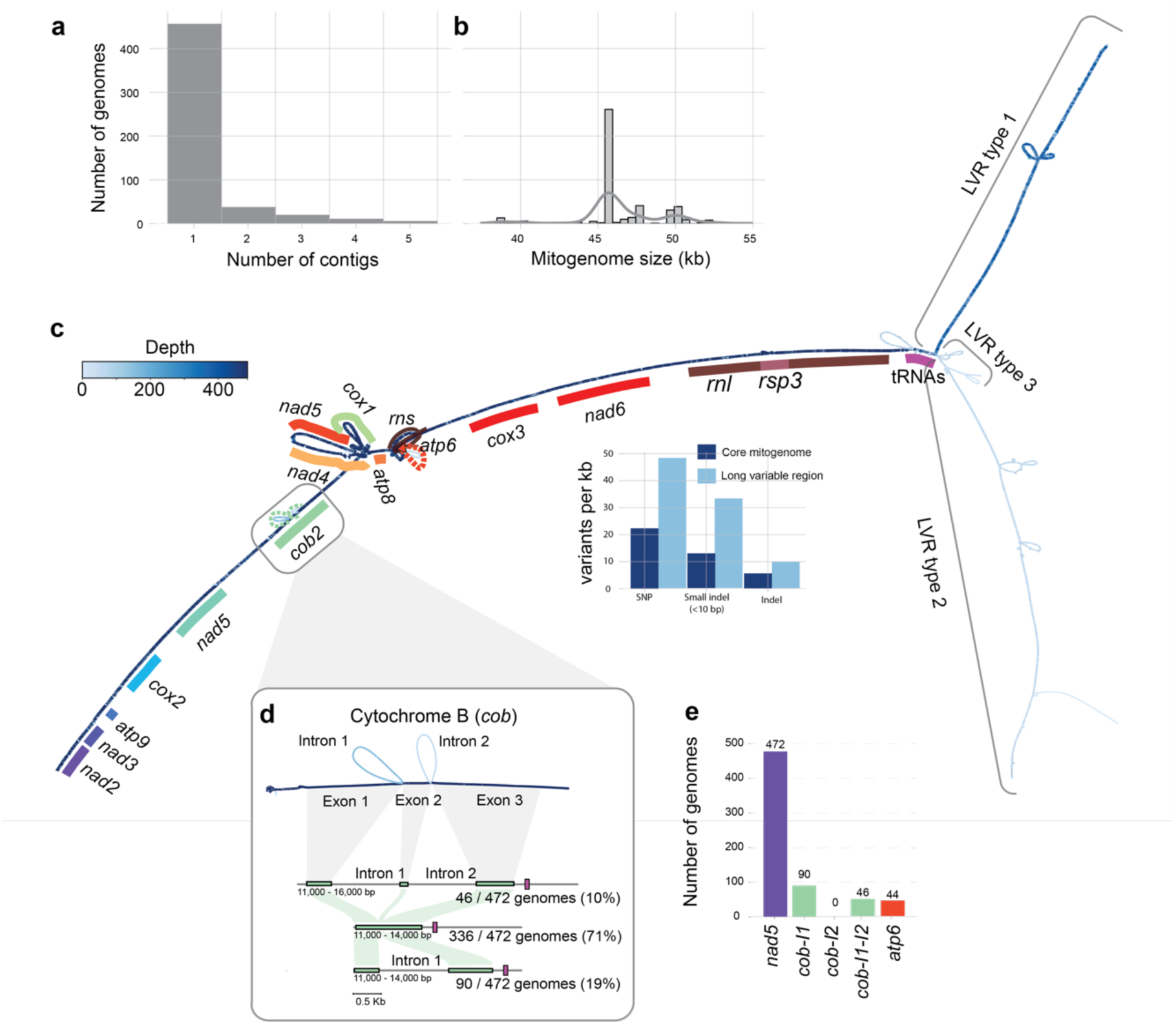
– Pan-mitogenome graph of *Fusarium oxysporum*. (**a**) The vast majority (472) of *F. oxysporum* mitogenomes were assembled into a single, circular contig. **(b)** The mitogenomes of *F. oxysporum* range in size between 38 – 55 kb. **(c)** The pan-mitogenome graph of *F. oxysporum* shows a highly conserved core region. In addition, the pan-mitogenome reveals three different long variable regions (LVRs) that are present in a subset of the mitogenomes. The dark blue colour indicates this region of the pangenome graph is present in all 472 mitogenome assemblies, while the lighter colours indicate that specific regions are present in fewer mitogenomes. **(d)** Pan-mitogenome graph uncovers variation in three out of four introns (seen as bubbles in the graph). The Cytochrome B (*cob*) gene has two different introns (*cob* I1 and I2), and 136 genomes contain at least one intron in *cob*. However, the pan-mitogenome graph shows there is no variation in the insertion site of the introns. **(e)** In total, four introns are found in the mitogenomes. All mitogenomes contain the intron in *nad5*, and thus this intron is not visible as an alternative path in the pangenome graph.

### Pan-mitogenome graph reveals highly conserved mitogenome sequences in *Fusarium oxysporum*

To compare the large collection of complete mitochondrial genomes, we constructed a pangenome variation graph using the Pan Genome Graph Builder (PGGB) (Fig. 2c). The pan-mitogenome graph has a length of 87,336 bp and is divided over 15,919 nodes connected by 22,498 edges. On average, a node is present in 189 mitochondrial genomes, and 1,974 nodes (27,552 bp) are present in all isolates (i.e. core), 12,206 nodes are accessory (46,887 bp), and the remaining 1739 nodes are unique to a single strain (12897 bp). To further quantify the variation between mitogenomes, we deconstructed sequence variants directly from the pangenome graph and identified a total of 4,961 variants (2,661 SNPs, 1,698 small indels (≦10 bp), and 602 larger indels (>10 bp) (Fig. 2b); no inversions or translocations were observed (Fig. 2c, Fig S1a). The pangenome graph shows only a single entangled region (Fig. 2c), in which some genomes traverse nodes more than once (Fig. S1b). This region localized at around 10 kb and contains *cox1*, *nad5*, *nad4*, *atp8*, and *atp6* (Fig. 2c). In this region almost all homing endonuclease motifs (HindII, CbaI, BamHI) are located, which occur in different copy numbers in the mitogenomes, and causes the ‘knotting’ of the graph. Importantly, the limited number of sequence variants as well as the lack of inversions and any other major structural variants demonstrate that the *F. oxysporum* mitochondrial genomes are largely conserved and syntenic, in line with the highly conserved gene order among Sordariomycetes (Fig. 1a).

### *Fusarium oxysporum* mitogenomes have few and conserved introns

Although the core mitogenome is conserved and syntenic between the 472 *F. oxysporum* mitogenomes, the core mitogenome encodes a variable number of introns, ranging from one intron in *nad5* (present in all 472 genomes) to four introns in *nad5*, *cob*, and *atp6* (present in 44 genomes) (Fig. 2a). Three of these introns are clearly visible as alternative paths in the pangenome graph, two in *cob* and one in *atp6* (Fig. 2c,d); the intron in *nad5* is present in all genomes and thus not visible as a variant. All four introns are self-splicing group I introns. Group I introns often encode homing endonuclease genes (HEGs) (Lang et al. 2007; Megarioti and Kouvelis 2020), which can be separated into different groups based on sequence domains. Three out of four introns (nad5, cob intron-II, and atp6) encode ‘LAGLIDADG’ domains, only the cob intron I encodes a ‘GIY’ domain. This is expected for fungal mitochondrial introns where most introns encode ‘LAGLIDADG’ domains (Megarioti and Kouvelis 2020).

Homing endonuclease genes in mitochondrial introns are mobile and can spread within and between genomes (Megarioti and Kouvelis 2020; Sellem and Belcour 1997). Using the location of the introns in the pangenome graph, we sought to identify different independent incursions of these introns into the mitogenomes; several intron incursions suggest high intron activity. The *cob* gene has an intron 392 bp after the start of the gene (intron I) and 488 bp after the start of the gene (intron II). We observed that a genome has either only intron I (19% of the genomes) or both intron I and II (10% of the genomes), yet we never observed intron II separately. The conserved location of these two introns indicates that these are the only intron incursions in cob. Importantly, the pan-mitogenome graph also does not uncover any sequence variants (represented as bubbles) within the intron, thus the sequence of the introns is highly conserved. The graph similarly shows that the introns in *atp6* and *nad5* are always located at the same location within the gene and no variants are found, corroborating previous analyses on a more limited set of mitogenomes (Brankovics et al. 2017) and highlighting that introns in *F. oxysporum* mitochondrial genome are conserved with no additional intron incursions. Interestingly, this contrasts to the intron expansions described in many other fungal mitogenomes (Fonseca et al. 2021; Wang et al. 2021), including those of *Fusarium graminearum* (Brankovics et al. 2018).

In many fungal mitogenomes, the abundance of introns correlates with the total mitogenome size (Fonseca et al., 2021; Wang et al., 2021, Fig. 1c). We similarly observed genome size variation in the 472 mitogenomes (Fig. 2b), yet due to the low number of introns in *F. oxysporum* mitogenomes, the introns at maximum describe up to ∼3,000 bp size variation. Consequently, we observed that the total genome size does not strongly correlate with the total number of introns (Pearson r. = 0.39, p < 0.05, Fig. S2), highlighting that different regions of the mitogenome must contribute the observed size variation.

### Variation between mitogenomes is caused by three different long variable regions unique to the *Fusarium* genus

Next to the conserved core region, the pan-mitogenome graph also clearly shows the presence of three different long variable regions (LVRs) (Fig. 2c, Fig. 3); the LVRs are visible in the pangenome graph, where the graph splits into three different paths. The three LVRs are localized between the *rnl* and *nad2* gene and flanked by tRNA clusters. These LVRs encode different open reading frames (ORFs) and vary largely in size (Fig. 3); LVR1 is approximately 10 kb, LVR2 18 kb, and LVR3 is the smallest with approximately 5 kb. The LVR size is strongly correlated with the size of the complete mitogenome (Pearson r = 0.89, p <0.05, Fig. S2), demonstrating that genome-size variation between *F. oxysporum* mitogenomes is driven by the type of LVR. Interestingly, previous reports based only on a small subset of *F. oxysporum* mitogenome report the same three long variable regions (Brankovics et al. 2017), and thus, unanticipatedly, the inclusion of hundreds of diverse mitogenomes does not reveal additional LVRs.

**Figure 3.**
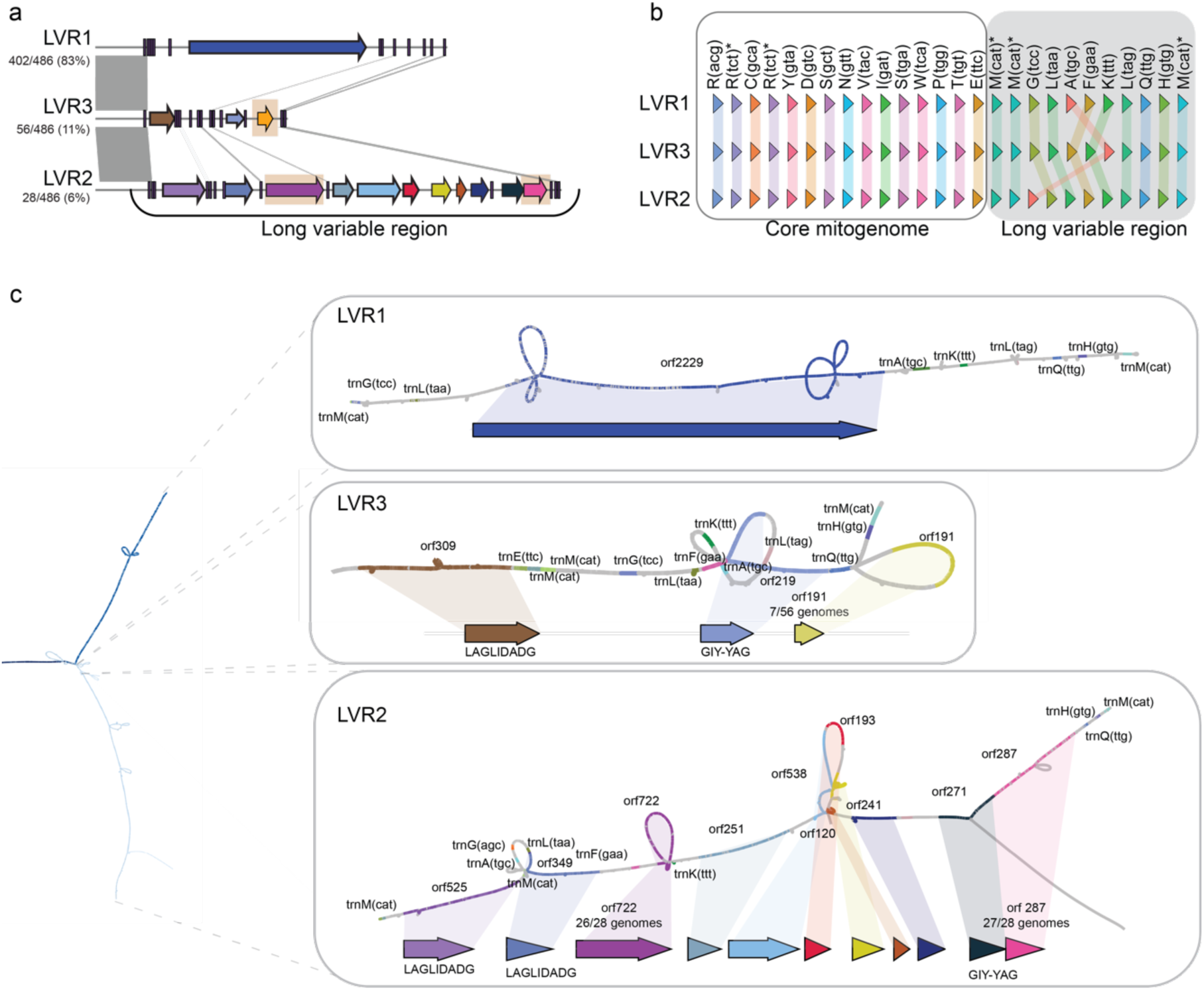
– The three long variable regions (LVRs) differ but variation within a LVR type is low. **a**) The three long variable regions (LVR) share little sequence similarity. Similar regions (indicated by grey lines) encompass only tRNA genes. Predicted open reading frames (ORFs) do not share sequence similarity. Orange boxes indicate ORFs that vary in presence/absence in the different mitogenomes. **b**) Comparison of the presence and position of tRNA genes in three mitogenomes with different LVR. All mitogenome contain similar tRNAs, both within the core mitogenome as well as within the LVRs. However, the order of tRNAs is specific for the different LVRs. **c**) The pan-mitogenome graph reveals limited variation in the different LVRs. Bubbles in LVR2 and 3 highlight presence/absence variation of ORFs (see also panel **a**).

The different LVRs have an unequal distribution between *F. oxysporum* strains. LVR1 is the most abundant and is detected in 83% of the mitogenomes, while LVR2 is found in only 6%, and LVR3 in 11% of the genomes (Fig. 2c, Fig. 3a). The GC-content of the LVRs (median 35.6%) differs from the median GC-content of the core mitogenome (31.0%). However, the GC content of the three LVRs varies as well. LVR1 has the highest GC content (35.7%), followed by LVR2 (32.4%) and LVR3 (31.9%). This shows that the higher median GC-content of the LVRs is mainly driven by the high GC-content of the most abundant LVR1. Interestingly, we observe a median GC-content of 27.5% in the coding regions and a GC-content of 34.3% in the intergenic regions. Is possible that the lower GC content in LVR2 and LVR3 is caused by the high abundance of ORFs in these LVRs. Importantly, the LVRs share no similarity to each other (Fig. 3a), this lack of homology as well as the absence of degenerated or chimeric LVRs suggests that the different LVR types are not derived from each other. The only similarity between the three LVRs is caused by sequence similarity between tRNAs (Fig. 3a,b). While the same tRNAs occur in all three LVRs, their order differs (Fig. 3b). In general, we find between 23 and 27 tRNAs, trnR(ttc) is found two times and trnM(cat) is triplicated. The trnM(cat) copies flank the LVRs, and all LVRs end with the same three tRNAs (Fig. 3b), and these tRNAs could therefore form putative recombination sites. Apart from the similar tRNAs, no small internal repeats are found in the LVRs.

The three distinct LVRs have been described in *F. oxysporum* (Brankovics et al. 2017), yet no biological function could be assigned nor has the origin of these regions been elucidated. The three types of LVRs encode different open reading frames (ORFs) without any similarity to each other (Fig 3) and also the number of ORFs per LVR varies. LVR1 contains only one ORF (ORF2290), and this ORF is present in all isolates encoding LVR1. In some genomes (76 out of 400 genomes) this ORF is truncated and split into several smaller ORFs. Most ORFs are found in LVR2, this region encodes nine ORFs of which seven are present in all isolates and two are variable (ORF722, absent in two genomes; ORF287, absent in one genome). LVR3 encodes three ORFs, of which one is present in only seven genomes. The variable ORFs are clearly present as bubbles in the graph (Fig. 3c), and the absences of two ORFs in LVR2 in only a low number of genomes suggests that these are recent, independent losses. On the other hand, ORF191 in LVR3 is present in only a low number of genomes, indicative of a recent gain. Notably, five of the ORFs encode homing endonuclease domains (Fig. 3c), however we did not observe any presence/absence variation of these in the ORFs. The other nine ORFs do not have any predicted function (table S4), and thus the function of the LVRs so far remains unelucidated.

To uncover the origin of the different LVRs, we sought to detect homologous regions in 706 diverse fungal mitogenomes (Fonseca et al. 2021) (Table S1). We observed that LVR1 is present in different *Fusarium* species, whereas LVR2 and LVR3 is only found in *F. oxysporum* (Fig. S4); outside the *Fusarium* genus, only small hits can be detected (Fig. S5). We also queried the non-redundant nucleotide database, excluding *Fusarium* species, and similarly only observed small (< 200 nt) matches (Table S5-7), suggesting that the most likely origin of these regions is within the *Fusarium* genus. Apart from the lack of homology to mitochondria, we also do not find any homology to nuclear elements both within and outside of the *Fusarium* genus.

### LVR types are incongruent with the nuclear phylogeny

Since shorter variants of the LVR, the absence of an entire LVR, or the presence of chimeric LVRs, might not be visible from the graph directly, we clustered the genomes based on the presence/absence pattern of the nodes in the pan-mitogenome graph (Fig. 4a). The clustering of the LVR reveals the presence of only three distinct LVRs types, in line with the observations from the pangenome graph (Fig. 2c, Fig. 3). To further analyse the various types of mitogenomes in *F. oxysporum*, we also clustered the complete pangenome graph. This revealed the same three clusters of mitochondrial genomes (Fig. 4a) and the separation into these three mitochondrial types is caused by the LVR, as the same three clusters are obtained by clustering solely the presence-absence of the LVR (adjusted rand index = 0.98, Fig. 4c). In contrast to the similarity between the complete graph and the LVRs, clustering based on the core mitogenome region uncovered four vastly different clusters (ARI = –0.02), suggesting that the observed variation between mitochondrial genomes is largely driven by LVRs and that the LVR evolves separately from the core mitogenome.

**Figure 4.**
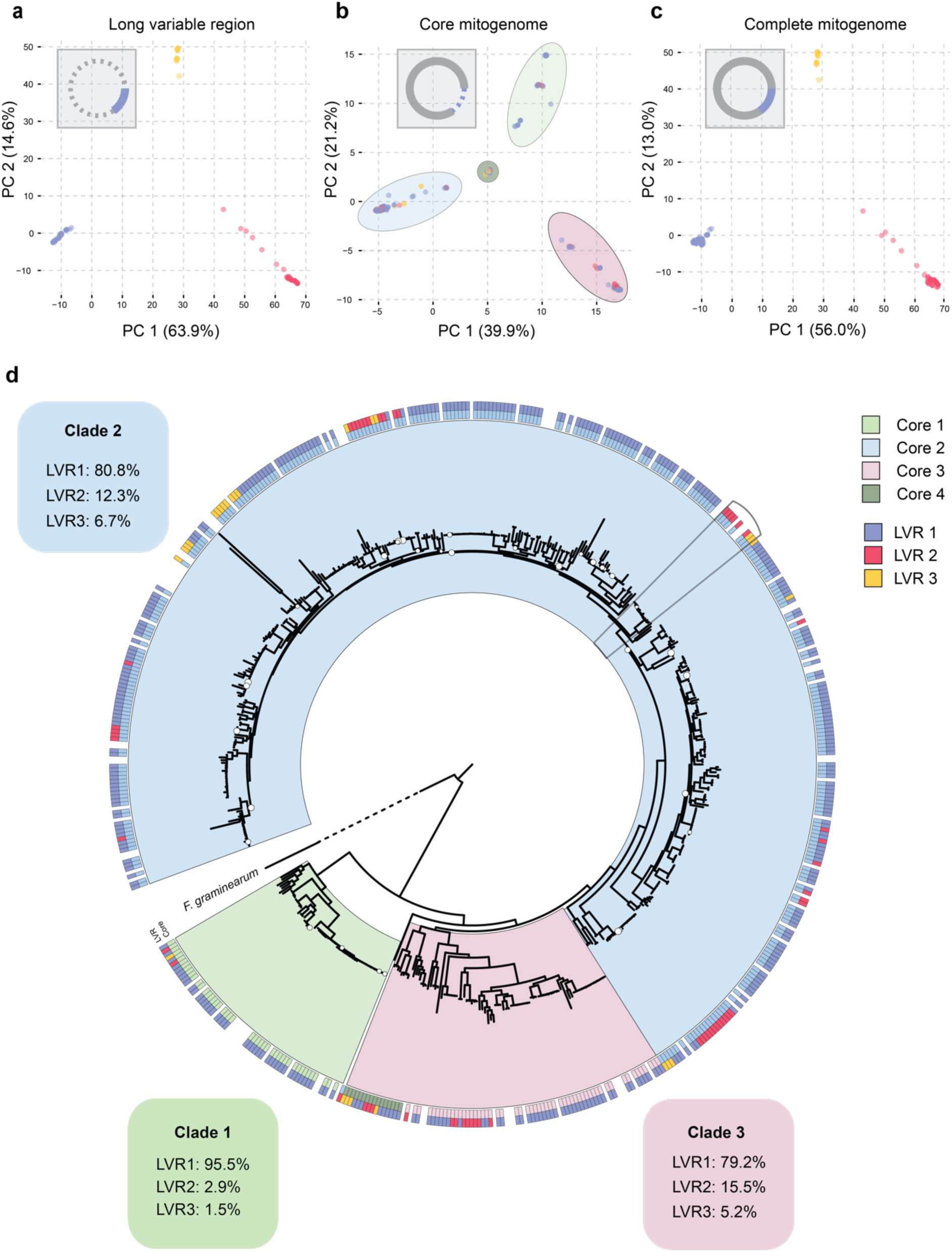
– Long variable regions are different from the core-mitogenome and do not correspond to lineages in the nuclear phylogeny. Principal component analysis (PCA) based on the presence / absence patterns of the nodes in **(a)** LVRs, **(b)** the core mitogenome, and **(c)** the complete pan-mitogenome graph. The core mitogenome results in four different clusters, the four groups are different from the groups obtained based on the complete pangenome graph (adjusted rand index = –0.02) and from the groups based on the long variable region (adjusted rand index = –0.02). **(d)** The nuclear phylogeny of 472 *F. oxysporum* assemblies is based on 4213 single-copy BUSCO genes and maximum-likelihood analysis (2,495,112 aa positions). The three well-recognized clades in *F. oxysporum* (O’Donnell et al. 1998) are clearly visible based on the nuclear phylogeny. The first outer ring indicates where the isolate clusters based on the core mitogenome. These core-mitogenome clusters as defined by the PCA (panel **b**) are similar to the taxonomic clades visible from the nuclear phylogeny. For one group of isolates in clade 3 the core mitogenomes cluster in the middle of the PCA (group core 4). The outer ring indicates the assigned clusters based on the presence of different LVRs (panel **a**), and these do not follow the nuclear phylogeny as all LVR types can be identified in all three *F. oxysporum* clades. The grey box highlights a clonal group that encodes two different LVRs.

To further trace the evolution LVRs in *F. oxysporum*, we compared the occurrence of LVRs and core-regions of the mitogenomes to a nuclear phylogeny that is based on nuclear-encoded, conserved, single-copy orthologs (Fig. 4d). The core mitogenomes can be separated into four groups based on the variants in the graph (PCA, Fig. 4b). The same four groups are also found based on a phylogenetic analysis using the 14 conserved core mitochondrial genes (Fig. S3). Nuclear clade 1 and 2 fully consist of only one core mitotype, however, clade 3 consist of core mitotype 3 and 4 (Fig. 4d, Fig. S3). Importantly, all genomes belonging to core mitotype 4 group together based on the nuclear (Fig. 4d) and core mitogenome phylogeny. This means that we observe additional variation in the core mitogenome, however the groups found based on the core mitogenome are in line with the nuclear phylogeny.

In contrast, the three different LVRs do not follow the nuclear phylogeny, and all three LVR-types occur in all three taxonomic clades (Fig. 4d). Interestingly, even genetically related F. oxysporum isolates can carry different LVRs. For example, two isolates in clade 2 have a highly similar core mitogenome (>99.9% identity, eleven SNPs, 14 small indels ≦ 10 bp, and four indels), yet they have different LVRs (LVR1 and LVR2) with only limited similarity around the tRNAs sequences (Fig 3b). Out of the 50 clonal groups in our dataset (branch length < 0.0001; on average five genomes per group, maximum of 35 genomes in the TR4 lineage), only one groups has more than one LVR (Fig. 4d). This indicates that it is possible for highly similar genomes, and even within a clonal group, to have different LVRs.

The incongruence between LVRs and the nuclear phylogeny might be due to horizontal transfer. In plants and fungi, horizontal transfer between mitochondria has been reported as well as the exchange of entire mitogenomes (Bergthorsson et al. 2004; Wu et al. 2015). The exchange between mitogenomes could result in heteroplasmy, i.e. the presence of two or more distinct mitochondrial populations within a single individual. To detect the occurrence of heteroplasmy, we compared the sequencing data of each of 472 isolates directly to the pangenome graph. We expected bi-allelic variants to indicate heteroplasmy, yet we did not observe any diploid variants in any of the isolates, neither in the core nor in the LVRs, indicating that heteroplasmy, if existent, is a very transient state in *F. oxysporum*. We then sought to experimentally examine if LVRs could be horizontally transferred between *F. oxysporum* stains by inducing a swap of LVRs between two related strains (strain NRRL36117 (LVR1), C135 (LVR2)). Although heterokaryons were successfully formed (Fig. S5), no exchange of mitochondria could be observed. Out of 47 spores derived from heterokaryotic mycelia, the nuclear genome and LVR of 44 spores was similar to NRRL36117 and only three spores were positive for C135 (Fig. S5). One of the spores remained a heterokaryon and was positive for both NRRL36117 and C135 primers (Table S3). This means we observed an unequal distribution of the two strains after heterokaryon formation, since most of the spores resemble NRRL36117. This is in line with previous reports that suggest that only one genome is maintained after heterokaryon formation (Shahi et al. 2016).

### Long variable regions are frequently exchanged

Possibly, the LVRs of mitochondria could be transferred between strains, yet the three different types can also be present in the ancestor. We expect that if all three LVRs are ancestral, their evolution would be in line with the nuclear genomes. We therefore constructed phylogenetic trees for the three LVRs and observed for LVR2 and LVR3 that genetically distant isolates can share the same or very similar LVR. For example, LVR3 from Fo6 (clade 1), Fo13 (clade 3), and Fo5 (clade 2) (all endophytes sampled from tomato plants in Australia) is highly similar (99.1 % identity, four small indels ≦10 bp), whereas the three genomes all belong to different taxonomic clades, and the core mitogenomes has 184 variants (84 SNPs, 73 small indels < 10bp, and 27 indels > 10 bp) as well as variation in intron presence / absence (Fig. 5c,d). We added an additional isolate from clade 2 carrying LVR3. This shows that the core mitogenomes are highly similar between the clade 2 isolates, yet LVR3 is only 96.1 % similar. Thus, LVRs from genetically distinct isolates, even belonging to a different phylogenetic clade, can be similar, while the core mitogenome shows variation between the strains. In total we observe four groups in the phylogeny of LVR2 and LVR3 where isolates from different clades cluster together based on the LVR. The unexpected similarity of LVRs between genetically distant strains can possibly be explained by the transfer of the LVR between isolates.

**Figure 5.**
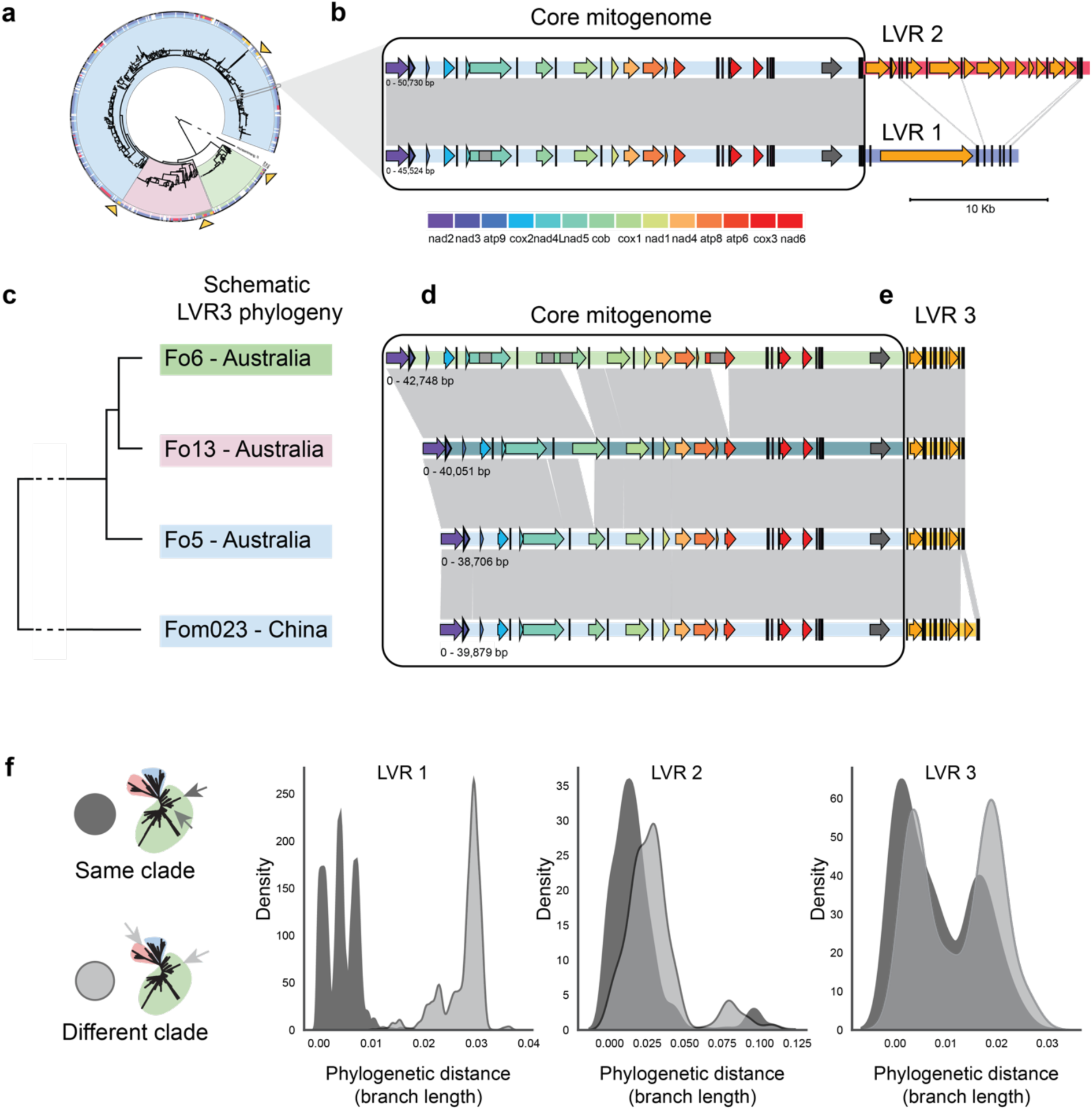
– LVRs are frequently exchanged between *F. oxysporum* isolates of different taxonomic clades. (**a**) Simplified nuclear phylogeny of *F. oxysporum* strains (see Fig. 4 for details), grey box indicates the strains compared in panel **b** and arrows indicate the strains compared in panel **c**. **(b)** Comparison of two closely related strains with different LVRs. Light blue region is the core mitogenome found in clade 2. The core mitogenome is highly similar, whereas LVR1 and LVR2 do not show any similarity (grey lines), apart from similar tRNA genes (black bars). **(c)** Schematic tree of LVR from four different strains belonging to different clades in the nuclear phylogeny (Fo6 clade 1, Fo13 clade 3, and Fo5 clade 2, indicated by orange arrows). **(d)** Comparison of the core mitogenome of three strains from a different clade carrying the same LVR3 variant, as well as a clade 2 strain carrying a different LVR3 variant (Fom023). The core mitogenome shows variation in intron content as well as some small intergenic variants. However, the LVR is highly similar (99.1%) between the isolates (grey lines) **(e)**. Fom023 encodes an extra open reading frame (orange arrow) in the LVR3. **(f)** Branch length between all possible pairs in each LVR phylogeny. For LVR1 the median branch length is higher between strains belonging to different clades (0.029) than for strains belonging to the same clade (0.004). For LVR 2 and LVR3, the median branch length between strains belonging to different clades (LVR2 = 0.027, LVR3 = 0.0136) is closer to the median branch length between strains belonging to the same clade (LVR2 = 0.0132, LVR3 = 0.006) and the distribution of branch lengths looks similar. This means that a LVR from isolates from different clades can be highly similar.

To further analyse the exchange of LVRs between different *F. oxysporum* clades, we directly compared the phylogenetic tree based on the LVRs to the phylogenetic tree based on the nuclear genome and quantified the phylogenetic distances (estimated by the branch length) between LVRs of isolates belonging to the same phylogenetic clade and between LVRs of isolates belonging to different clades. Interestingly, the phylogeny of LVR1 shows three distinct clusters similar to the three clades found in the nuclear phylogeny (Fig. S6). Thus, isolates with LVR1 belonging to the same clade typically carry a similar LVR, separated by a low phylogenetic difference (median branch length = 0.004), whereas isolates belonging to different clades have more dissimilar LVR1s (median branch length = 0.029, p = 0.00, Fig. 5f). This suggests that LVR1 evolves in line with the nuclear genome and that LVR1 was likely present in the last common ancestor of *F. oxysporum*. However, the pattern for LVR2 and LVR3 is remarkably different. Here, the similarity of LVRs between isolates belonging to the same taxonomic clade (LVR2: mean branch length = 0.0132, LVR3: mean branch length = 0.006) is close to the similarity between LVRs of isolates belonging to different taxonomic clades (LVR2: mean branch length = 0.027, LVR3: mean branch length = 0.0136), which is in line with the highly similar LVR3 that can be found in three isolates from different taxonomic clades (Fig. 5c-e, Fig. S8). These results not only suggest that LVRs can be exchanged between strains, but that this exchange can occur between genetically distant strains that belong to different taxonomic clades.

## Discussion

The evolutionary history of mitogenomes can provide direct insights into the evolutionary history of their host species (Christinaki et al. 2022; Ingman et al. 2000). Pangenome graphs recently emerged as a novel framework to facilitate analyses of large genomic datasets and are a useful tool to uncover novel biological insights (Li et al. 2022; Wang et al. 2022b). We here constructed, to our knowledge, the first pan-mitogenome of a filamentous fungus, based on hundreds of genetically diverse *F. oxysporum* isolates. Many members of the large and diverse *F. oxysporum* species complex are major plant pathogens, and for a long time these have been considered to only reproduce clonally. However, recently signatures of sexual recombination have been observed, indicating the presence of a sexual cycle in this presumed asexual fungus (Fayyaz et al. 2023; McTaggart et al. 2021). Previously analyses of the mitogenome of *F. oxysporum* provided initial evidence for mitochondrial recombination (Brankovics et al. 2017), and we here sought to further elucidate these mitochondrial dynamics and the patterns of recombination. Our pan-mitogenome graph uncovers highly conserved and syntenic core mitogenomes. In contrast, we could clearly distinguish three distinct types of long variable regions (LVRs), which explain most of the mitochondrial variation in *F. oxysporum*. Whereas the core mitogenome largely evolves in line with the nuclear genome, the LVRs are incongruent with the nuclear phylogeny. Importantly, we found highly similar LVRs between genetically distant isolates, even between isolates from a different phylogenetic clade, which shows that genetic exchange can occur between distantly related *F. oxysporum* isolates, suggesting that distant taxonomic clades, contrary to previous expectations, are not isolated. This seemingly unrestricted capacity to recombine has significant implications in the evolution of *Fusarium*, for instance in their ability to horizontally exchange accessory chromosomes.

LVRs have been discovered previously in *Fusarium* and are found to encode open reading frames (ORFs) of unknown function (Al-Reedy et al. 2012; Brankovics et al. 2017). Similarly, ORFs of unknown function are described, for example in other fungal species (Jelen et al. 2016) and bivalves (Milani et al. 2013). In the latter case, the unknown ORFs seems to be derived from viral origin and play a role in the inheritance of mitochondria (Milani et al. 2013). However, these described ORFs are shorter than the long ORF found in LVR 1 in the *Fusarium* genus and the origin and function of these ORFs remain unclear. Notably, we here observed only three different types of LVRs in the *F. oxysporum* pan-mitogenome and even though we included nearly 500 diverse isolates, no degenerated or chimeric variants could be observed. Previous studies suggested that LVRs evolve similar to intergenic regions. We similarly determined that LVRs have a higher density of variants per kb than the core mitogenome. The lack of homologous sequences in other species hampers our attempts to further elucidate the origin of the LVR. The abundance of LVR1, together with the presence of LVR1 in other *Fusarium* species members and the similarity to the nuclear phylogeny, suggests that this LVR is most likely ancestral. However, we did not find similarity between LVR1, LVR2, and LVR3, showing that these types do not originate from recombination with LVR1 or through degeneration. Interestingly, we observe highly similar tRNA genes in the LVRs. This might suggest that the three LVRs originate from the same ancestral sequence and that the different LVRs evolved by integration of homing endonucleases or other mobile elements. However, we observe little variation within an LVR type and no presence absence variation of the homing endonuclease, suggesting these are not active. Therefore, we do not think that this hypothesis fits our observations. Importantly, LVR2 and LVR3 do not follow the nuclear phylogeny, and we hypothesize that the LVRs are exchanged through recombination between two mitochondria encoding a different LVR. It is conceivable that the similar tRNA genes that flank the LVRs form recombination sites and the clustering of these tRNAs at the flank of the LVRs drives the observed pattern of recombination (Fritsch et al. 2014; Pratt-Hyatt et al. 2006). Notably, while frequent recombination of LVRs between isolates seems to occur, we did not detect any signature of recombination in the core genome. It is, however, possible that recombination in the core region only occurs between highly similar isolates and therefore, there are no visible signs of recombination. This apparent lack of recombination in the core mitogenome is in line with the highly conserved gene order among *F. oxysporum* species and even among the class of Sordariomycetes, as gene order shuffling is the result of recombination (Aguileta et al. 2014). Alternatively, selection against additional recombination sites that affect the core mitogenome can result in a lack of observed recombination.

*F. oxysporum* strains belonging to different taxonomic clades can carry surprisingly similar LVRs, which suggest that the mitochondria of these isolates can recombine regardless of the extensive genetic differentiation between the nuclear genomes. Exchange of LVRs between genetically distinct isolates suggests the presence of mitochondrial recombination between different taxonomic clades. This is a significant finding as the prevailing thought is that mitochondrial recombination does not occur between taxonomic clades (Brankovics et al. 2017; McTaggart et al. 2021). To further interpret the incongruent phylogeny between the LVR and the nuclear genome, it is imperative to understand the mode of mitochondrial inheritance. In fungi, both uniparental and biparental mitochondrial inheritance has been described (Barr et al. 2005), and recombination of mitochondria has been observed. In yeast, for example, biparental inheritance results in short-lived heteroplasmy, during which mitochondrial recombination can occur (Wu et al. 2015). Although *F. oxysporum* has for long been considered to lack a sexual cycle, and thus the mitochondria should be inherited clonally, conidial anastomosis in filamentous fungi allow for the fusion of cells of distantly related isolates (Read et al. 2012; Roca et al. 2003). In this process, the cytoplasm of cells is connected, which enables the movement of mitochondria (Kurian et al. 2018; Mehta and Baghela 2021; Roca et al. 2003; Vangalis et al. 2021). Upon fusion, only one nucleus is maintained in *F. oxysporum* (Kurian et al. 2018), and it has been indicated in four strains that the mitotype of the acceptor is maintained upon horizontal transfer events (van Dam et al. 2017). This is in line with our own observations that suggest that after heterokaryon formation, only one nucleus and one mitogenome is maintained. Our data suggest that, even though we observed patterns of frequent recombination between mitochondria of extant strains, this is likely an infrequent event in nature. Notably, we observed exchange between strains that were sampled from the same geographic location or similar environment, as physical proximity of isolate is a prerequisite of genetic exchange. However, knowledge on mitochondrial fusion, fission, and recombination upon heterokaryon formation remains limited.

In fungi, horizontal gene and chromosome transfer occurs (Habig et al. 2021; Mehrabi et al. 2011). Extensive horizontal chromosome transfer has been observed in *F. oxysporum*, even between distantly related strains that are not vegetatively compatible, which means the strains cannot form a viable heterokaryon (Fokkens et al. 2018; Shahi et al. 2016; Vlaardingerbroek et al. 2016). The horizontal transfer of nuclear DNA over large genetic distances might occur parallel to the exchange of mitochondrial LVRs over large genetic distances. Yet, the exchange of chromosomes in *F. oxysporum* has only been observed between strains belonging to the same clade (Fokkens et al. 2018; Shahi et al. 2016; Vlaardingerbroek et al. 2016), whereas we observe recombination of mitochondria between strains belonging to different clades. This suggests that exchange of accessory chromosomes over large genetic distances might be possible, and that the clades are not genetically isolated as has been previously thought (Brankovics et al. 2017; McTaggart et al. 2021). It is important to note, however, that it is currently unclear how the dynamics of the mitogenome relate to nuclear genomes and, to our knowledge, the relationship between mitochondrial transfer and horizontal gene transfer has not been studied. Examples of mitochondrial exchange can form an excellent starting point to relate the exchange of mitochondrial material to the exchange between nuclear genomes.

The transfer of mitochondria between cells has been observed in various other species. For example, mitochondria can be exchanged between human tissues (Borcherding and Brestoff 2023) and between plant cells, for instance, upon grafting (Gurdon et al. 2016). Transfer of mitochondria enables mitochondrial recombination between varying mitogenomes. Extensive horizontal transfer of mitochondria has been reported in the basal angiosperm *Amborella*, where the mitochondrial DNA contains material from great phylogenetic distances like mosses green algae but also other angiosperms (Bergthorsson et al. 2004; Rice et al. 2013). However, it is currently unknown what mechanisms enable these mitochondrial transfers over large evolutionary distances.

Recent developments in mitogenome analyses have uncovered novel insights in mitochondrial biology, and the remarkable evolutionary dynamics of mitogenomes are becoming clear (Burger et al. 2003; Gurdon et al. 2016; Rice et al. 2013). This raises the questions how this genetic variation in mitochondrial genomes is established and how the evolutionary dynamic of mitochondria links to the biology and the evolution of their host species. We here demonstrate, by using *F. oxysporum* mitogenomes as an example, that pan-mitogenomes provide an important framework to elucidate mitochondrial genome evolution not only in fungi but also in other eukaryotes, which can ultimately help to understand their evolution.

## Materials and Methods

### Mitogenome assemblies

To determine the gene-order and intron content in fungal mitochondria, we downloaded 706 publicly available fungal mitogenomes (Fonseca et al. 2021). These genomes have been annotated using MFannot followed by manual curation (Fonseca et al. 2021).

To study the evolution of *Fusarium oxysporum* mitogenomes, we downloaded all available 813 short-read dataset from the NCBI short read archive (SRA) on September 11^th^, 2022 (Table S2). Additionally, we added 70 *Fusarium oxysporum* strains infecting banana (van Westerhoven et al. 2024). Because all data is obtained from different sources, we checked the dataset to make sure the short-read data is correctly labeled and of sufficient quality. We filtered the dataset on BUSCO completeness (version 5.3.1, hypocreales_odb10), selecting isolates containing more than 90% of the single copy BUSCO genes. Based on 4213 BUSCO genes we estimated the maximum likelihood phylogeny using FastTree version 2.1.11 (Price et al. 2010). We subsequently removed all isolates that were not associated to *Fusarium oxysporum* (details for data acquisition and filtering, SI methods).

### Assembly and annotation

We assembled all *Fusarium oxysporum* short reads using GrabB version 1 (Brankovics et al. 2016), providing six complete *Fusarium oxysporum* mitogenomes from the NCBI database (NCBI accessions: MT010933.1, KR952337.1, AY945289.1, MF155191.1, MT269799.1, and MT010933.1) to bait mitochondrial reads from the short-read data. To ensure high-quality and contiguous mitogenome assemblies, we continued only with the assemblies that were successfully assembled into a single contig.

The assembled mitogenomes were subsequently annotated using MFannot version 3.0, specifying the translation frame table 4 and allowing for small motif and homing endonuclease domain annotation (Lang et al. 2023). In addition to known mitochondrial genes, many ORFs were annotated per genome. To identified homologous ORFs in the different annotations, we clustered ORFs with CD-hit version 4.8.1 (Li and Godzik 2006), with sequence identity of 0.8 and an alignment coverage of the shortest sequence of 0.8. Protein domains in these ORFs were identified using Interproscan version 5.67 (Quevillon et al. 2005).

### Graph construction

To build the pangenome variation graph, we used pangenome graph builder (PGGB) version 0.3.0 (Garrison et al. 2023). The segment size (-s) was set to 3000 bp and the percent identity was calculated using mash triangle (Ondov et al. 2016), resulting in a percent identity (-p) of 97.523., asm10. The graph was visualized using Bandage version 0.9.0 (Wick et al. 2015). We further analyzed the graph using odgi version 0.8.2 (Guarracino et al. 2022), and general graph statistics were obtained using odgi stats. To annotate the graph, we provided the annotation of the *F. oxysporum* stains TR4 II5 mitogenome to odgi position, the resulting csv file was used to extract loci and to determine the gene locations in the graph. To determine the variants in the graph, we used the deconstruct command from vg-tools version 1.49.0 (Garrison et al. 2018) specifying flag –a to get nested variants, to create a vcf file of the variants in the graph.

To analyze potential heteroplasmy of mitogenomes in *Fusarium oxysporum*, we aligned all unassembled short reads to the graph using vg giraffe version 1.49.0 (Sirén et al. 2021) and obtained variants using vg call (Hickey et al. 2020) with the diploid flag. The number of diploid variants were used to estimate the presence of heteroplasmy.

### Analysis of LVRs

To analyze the different types of mitogenomes the graph, we obtained the node presence absence based on odgi matrix. This matrix was used for a principal component analysis using scikit learn version 0.24.2 (Pedregosa et al. 2011).

To construct the phylogeny of the LVRs, we extracted the LVR sequence from all mitogenomes. For each genome, we determined the genomic coordinates belonging to the LVR and extracted the fasta files using bedtools getfasta (Quinlan and Hall 2010). We aligned the LVR sequences using mafft version 7.520 (Katoh et al. 2002) and used these alignment to construct a phylogeny using RAxML version 1.2.0 (Stamatakis 2014) with the model ‘GTR+G’. We calculated the pairwise branch length between each pair of isolates in the LVR phylogeny using Biopython version 1.79 (Cock et al. 2009) and grouped the pairs in to two categories; either both genomes belong to the same clade or belonging to a different clade. To plot the similarity between mitogenomes we used Pygenomeviz version 0.4.4 (Shimoyama 2022). The genome alignments were calculated using mummer version 4 (Marçais et al. 2018).

We searched the presence of the three LVRs in the 706 fungal mitogenomes (Table S1), using one representative of each LVR (LVR1; M1, LVR2; NRRL36117; LVR3; SRR10428567 – Fo6) as a query for BLAST. To find potential hits with isolates not included in the set of fungal mitogenomes, the LVRs were searched against the BLAST database (accessed 15-05-2024), excluding hits with members of the *Fusarium* genus.

We further assayed the behavior of mitochondria upon heterokaryon formation of genetically related strains carrying different LVRs. To test if the mitotypes can be swapped between isolates, we created single-spore isolates from the heterokaryon and tested the LVR type present in this isolate (for details, see SI methods; Table S6).

## Data accesses

All mitogenome assemblies and gene annotations are available on Figshare (10.6084/m9.figshare.26063593). The scripts used in this study are available on Github (https://github.com/Anouk-vw/pan-mitogenome_FOSC).

## Acknowledgements

A.C.W, J.D, and G.H.J.K were supported by a grant from the Bill and Melinda Gates Foundation to the International Institute of Tropical Agriculture, Agreement No – AG-5797. The funders had no role in study design, data collection and analysis, decision to publish, or preparation of the manuscript. We thank Like Fokkens for her feedback on the manuscript.

## Author contributions

A.C.W performed analysis and data analyses, and wrote the draft manuscript, with support of M.F.S. J.D conducted experiments. L.A.P, K.W and J.B were involved in the data analysis and interpretation. All authors contributed to editing of the manuscript. G.H.J.K and M.F.S supervised the project.

