## Supplementary Material & Methods for "Frequent genetic exchanges revealed by a pan-mitogenome graph of a fungal plant pathogen"

**Fungal mitogenomes**

To determine the patterns of gene-order and intron content in fungal mitochondria, we downloaded 706 fungal mitogenomes (Table S1, (Fonseca et al., 2021). These genomes have been previously annotated using MFannot, and were subsequently manually curated (Fonseca et al., 2021). We parsed this annotation to obtain the order of 14 fungal core genes. Fungal strains were ordered based on the NCBI taxonomy information, parsing the NCBI identifier using ete3 (Huerta-Cepas et al., 2016). Per genome, we counted the number of introns and correlated their abundance to the mitochondrial genome size using scipy version 1.5.3 (Virtanen et al., 2020).

***Fusarium oxysporum* short-read archive data**

To generate a pan-mitogenome including all publicly available sequencing data of *Fusarium oxysporum*, we downloaded all available 813 short-read dataset from the NCBI short-read archive (SRA) on September 11^th^, 2022. Only data from whole-genome sequencing efforts were included and we applied the following search terms “*("Fusarium oxysporum"[Organism] AND ("biomol dna"[Properties] AND "strategy wgs"[Properties] AND "library layout paired"[Properties] AND "platform illumina"[Properties] AND "filetype fastq"[Properties])”*. Additionally, we added a collection of 70 *Fusarium oxysporum* strains infecting banana (van Westerhoven et al., 2024). Because the data has been obtained from different sources, we aimed to make sure the short-read data is correctly labeled and of sufficient quality. We therefore assembled the nuclear genomes using spades version 3.13.0 (Bankevich et al., 2012) and ran BUSCO version 5.1.3 (Simão et al., 2015) with hypocreales_odb10 dataset to analyze the presence of conserved single-copy orthologs in the whole-genome nuclear assemblies. Genomes containing less than 90% BUSCO genes were excluded from further downstream analysis, as these genomes are likely incomplete, contaminated, or do not belong to the order of Hypocreales. This resulted in 623 assembled genomes.

Next to serving as a proxy for genome completeness, the BUSCO genes were also used to construct an approximate-maximum likelihood phylogeny of the here analyzed nuclear genomes. We collected 4213 BUSCO genes present in at least 95% of all 623 genomes, as well as the BUSCO genes present in *Fusarium graminearum* to represent the outgroup. The amino acid sequences for each BUSCO gene was aligned using muscle version 5.1 (Edgar, 2004) and subsequently concatenated, resulting in an alignment of 2,495,112 amino acid position. FastTree version 2.1.11 was used to construct the phylogeny, and the tree was visualized using iTol (Letunic & Bork, 2021). We subsequently removed genomes that cluster together with the outgroup, as we reasoned that these isolates do not belong to the *Fusarium oxysporum* species complex.

**Mitochondria swapping experiment**

To determine the presence of heterokaryon formation, *nit* mutants were generated for *Fusarium oxysporum* strains C135 and NRRL36117 (Parte et al., 2023), as described previously (Puhalla, 1985). In brief, agar plugs were placed on PDA plates supplemented with 1.5% KClO_3_. Plates were monitored for fast growing sectors for 14 days. Fast growing sectors were isolated to fresh MMA plates (Puhalla, 1985) to check for thin mycelial growth indicative of mutations in nitrate utilization. Collected *nit* mutants were paired on MMA with other mutants of the same strain to obtain at least two compatible mutants for both C135 and NRRL36117. Complementation was noted as the formation of dense hyphal growth starting from the hyphal contact point of two mutants. To test if C135 and NRRL36117 are part of the same VCG, pairings were made on MMA plates using two compatible mutants from both strains.

To test for the potential transfer of mitochondria after heterokaryon formation, agar plugs of compatible nit mutants of C135 and NRRL36117 were paired on MMA plates. Dense aerial hyphae were sampled for PCR testing using primers specific for the different core and mitochondrial genomes (Table S3). Single-spored colonies were generated from the aerial hyphae from the heterokaryotic aerial hyphae and tested for potential mitochondrial transfer using the same primers.
