## Supplementary Figures for "Frequent genetic exchanges revealed by a pan-mitogenome graph of a fungal plant pathogen"

- **Table S1**: Overview of 706 fungal mitogenomes
- **Table S2**: *Fusarium oxysporum* short read datasets included in this study
- **Table S3**: PCR primers used to validate the identity of the nuclear genome and the mitochondrial long variable region
- **Table S4**: BLAST hits of LVR 1 against the nucleotide database excluding members of the Fusarium genus
- **Table S5**: BLAST hits of LVR 2 against the nucleotide database excluding members of the Fusarium genus
- **Table S6**: BLAST hits of LVR 3 against the nucleotide database excluding members of the Fusarium genus
- **Figure S1**: Visualization of nodes that are present or absent in the *Fusarium oxysporum* pan-mitogenome sequence graph. Each row corresponds to a path (i.e., mitogenome) in the pangenome graph showing nodes (sequences) as colored rectangles. (**a**) Nodes are colored by the inversion rate per node. Black nodes indicate no inversion and red nodes would indicate inversions. Importantly, all nodes in the pan-mitogenome graph are colored black, and thus no inversions are present. (**b**) Nodes are colored by the coverage per node. Red nodes correspond to highly covered repetitive nodes. (**c**) Nodes are colored by the nucleotide position in the mitogenome. Light grey nodes correspond to the starting position (*nad2*), darker colors indicate position further in the mitogenome, which demonstrates that all mitogenomes follow a similar conserved order.
- **Figure S2** – Correlation between mitogenome size and (**a**) number of introns and (**b**) length of the long variable region (LVR). The mitogenome size is only weakly correlated to the number of introns (Pearson r = 0.37, p < 0.05) but a strong correlation is observed between the mitogenome size and the length of the LVR (Pearson r = 0.88, p < 0.05).
- **Figure S3** – Comparison of the three long variable regions (LVR) identified in the pan-mitogenome graph. Importantly, the three LVRs do not show any sequence similarity, except for some of the tRNA genes (purple boxes). The LVRs vary in size as well as in the abundance and number of predicted open reading frames (orf) (orange arrows).
- **Figure S4 –** BLAST search of the three long variable regions (**a)** LVR1, (**b)** LVR2, (**c)** LVR3) against 706 fungal mitogenomes (supplementary table 2), indicating that the LVRs are only found in members of the Fusarium genus, outside the Fusarium genus very short hits are found. LVR 1 is found outside of Fusarium oxysporum, whereas LVR 2 and 3 are soley found in members of the *Fusarium oxysporum* species compex. The color indicates the percent identity per genomic region.
- **Figure S5** – PCR identification of the nuclear genome and the mitochondrial long variable region (LVR) upon heterokaryon formation of *Fusarium oxysporum* strain NRRL36117 (LVR 2) and C135 (LVR 1). (**a**) Growth of the C135 and NRRL36117 *nit*-mutants. Upon heterokaryon formation growth of the mutants is restored (red circle). The heterokaryon was sampled and diluted to obtain single spore colonies. 47 colonies have been screen by PCR for **(b)** the C135 nuclear genome primer, **(c)** LVR 1, **(d)** NRRL 36117 nuclear genome and **(e)** the LVR 2 primer. Yellow boxes indicate three spores that are positive for the C135 nuclear genomes and the corresponding LVR1. One out of 47 colonies is still a heterokaryon (purple box) and positive for the NRRL36117 and C135 nuclear genome as well as for both LVRs. No exchange of LVRs is observed upon heterokaryon formation in this experiment.
- **Figure S6** – Maximum likelihood phylogeny of long variable region 1 (LVR1). Labels are colored according to the nuclear phylogeny. Based on the LVR1 phylogeny, isolates belonging to the same clade group together, suggesting that LVR1 largely follows the nuclear phylogeny.
- **Figure S7** - Maximum likelihood phylogeny of long variable region 2 (LVR2). Labels are colored according to the nuclear phylogeny. The phylogeny shows that isolates from different clades can share a highly similar LVR2. For example, R1B (Clade 1) groups together with other genomes from clade 2.
- **Figure S8** - Maximum likelihood phylogeny of long variable region 3 (LVR3). Labels are colored according to the nuclear phylogeny. Isolates belonging to a different clade can have a highly similar LVR3. For example, three genomes (SRR10428567; Fo6, SRR10428572; Fo13, SRR10428602; Fo5) of three different clades group together on one branch.

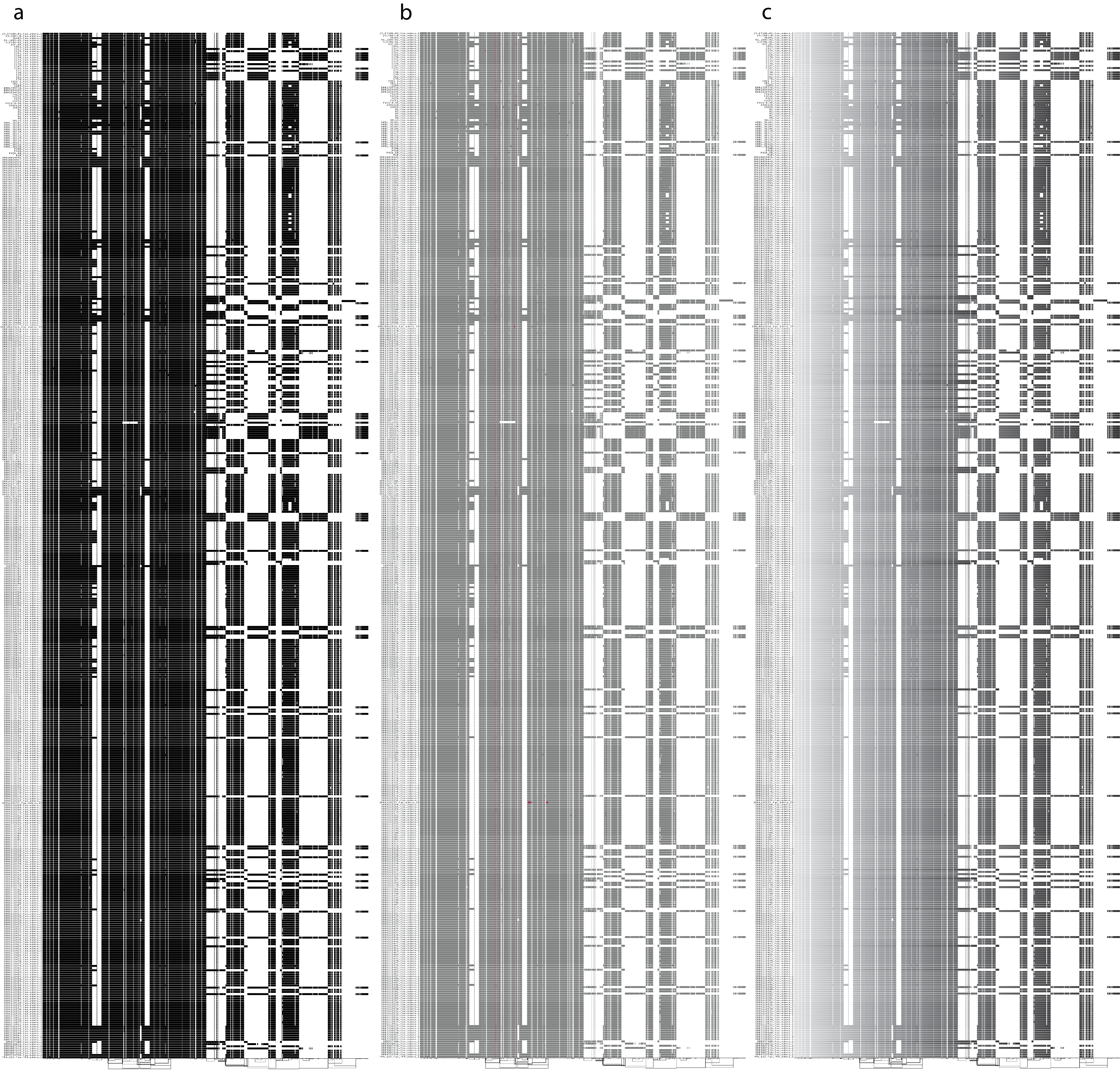

**Figure S1** – Visualization of nodes that are present or absent in the *Fusarium oxysporum* pan-mitogenome sequence graph. Each row corresponds to a path (i.e., mitogenome) in the pangenome graph showing nodes (sequences) as colored rectangles. (**a**) Nodes are colored by the inversion rate per node. Black nodes indicate no inversion and red nodes would indicate inversions. Importantly, all nodes in the pan-mitogenome graph are colored black, and thus no inversions are present. (**b**) Nodes are colored by the coverage per node. Red nodes correspond to highly covered repetitive nodes. (**c**) Nodes are colored by the nucleotide position in the mitogenome. Light grey nodes correspond to the starting position (*nad2*), darker colors indicate position further in the mitogenome, which demonstrates that all mitogenomes follow a similar conserved order.

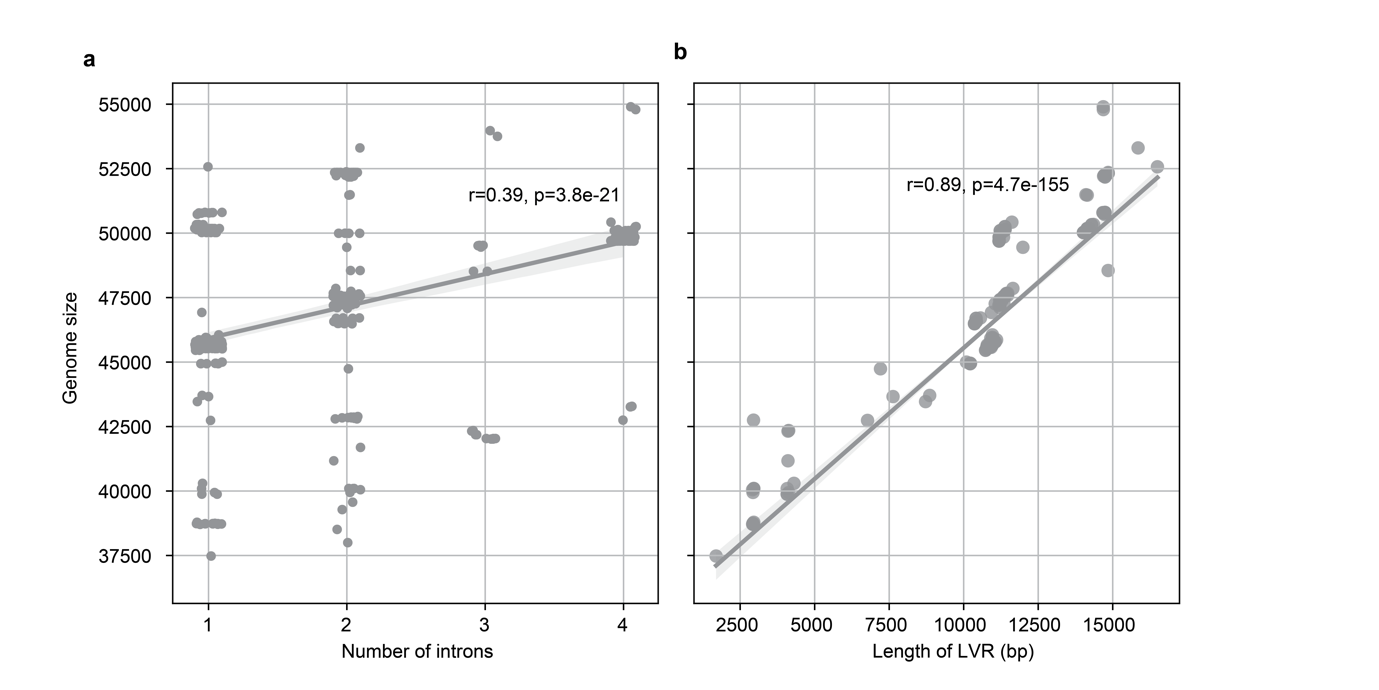

**Figure S2** – Correlation between mitogenome size and (**a**) number of introns and (**b**) length of the long variable region (LVR). The mitogenome size is only weakly correlated to the number of introns (Pearson r = 0.39, p < 0.05) but a strong correlation is observed between the mitogenome size and the length of the LVR (Pearson r = 0.89, p < 0.05).

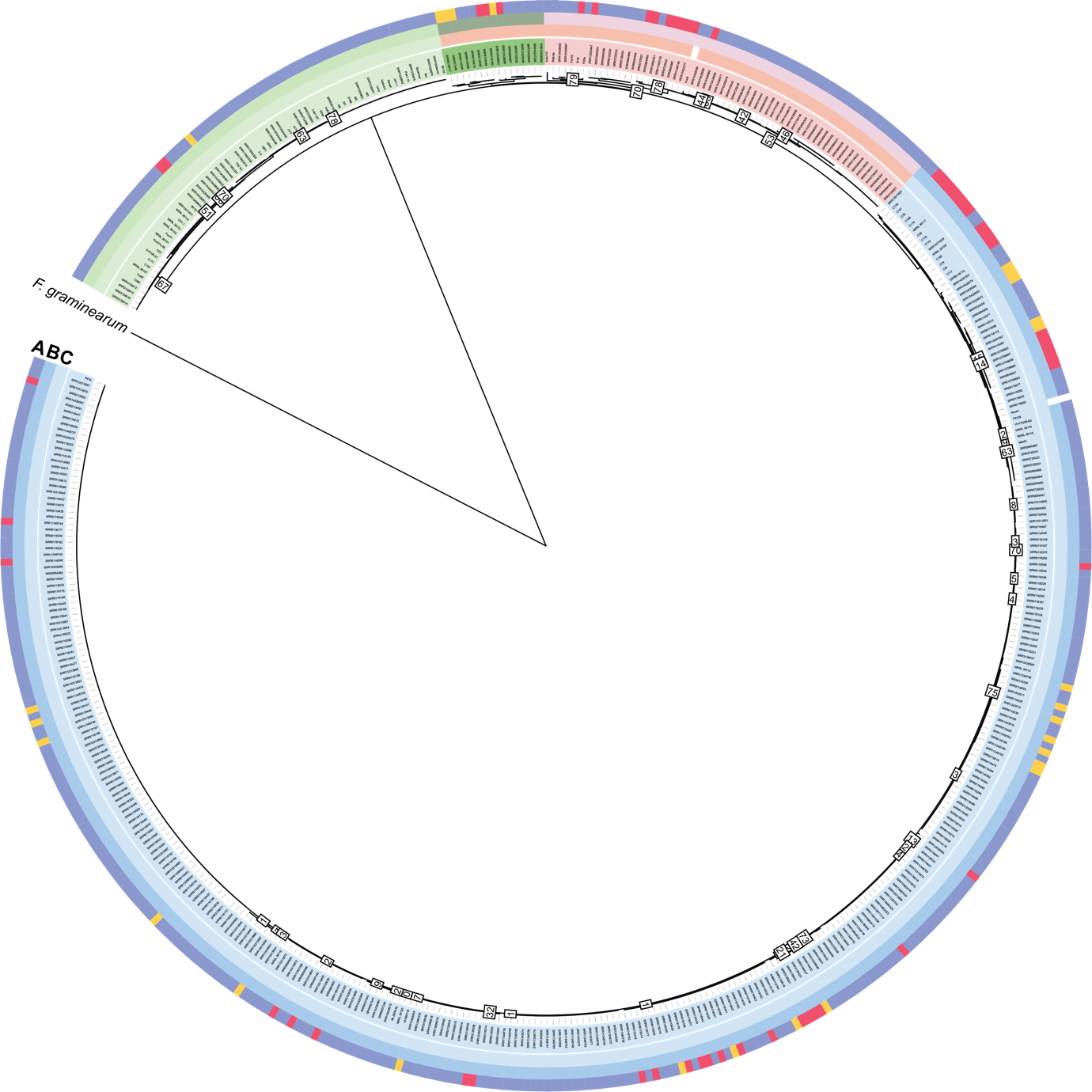

**Figure S3** –Phylogenetic tree based on 14 conserved mitochondrial genes. Bootstrap values are indicated for branches with a bootstrap < 70. Colors indicate a) Long variable region associated with the isolates b) core-mitogenome type determined based on the PCA. c) clade in the nuclear phylogeny. Leaf labels are colored according to the clades in this phylogenetic tree. Grey box indicate the isolates that carry core-mitogenome type 4. These isolates belong to clade 3 based on the nuclear phylogeny, but form a separate group in the PCA and in the core mitogenome phylogeny.

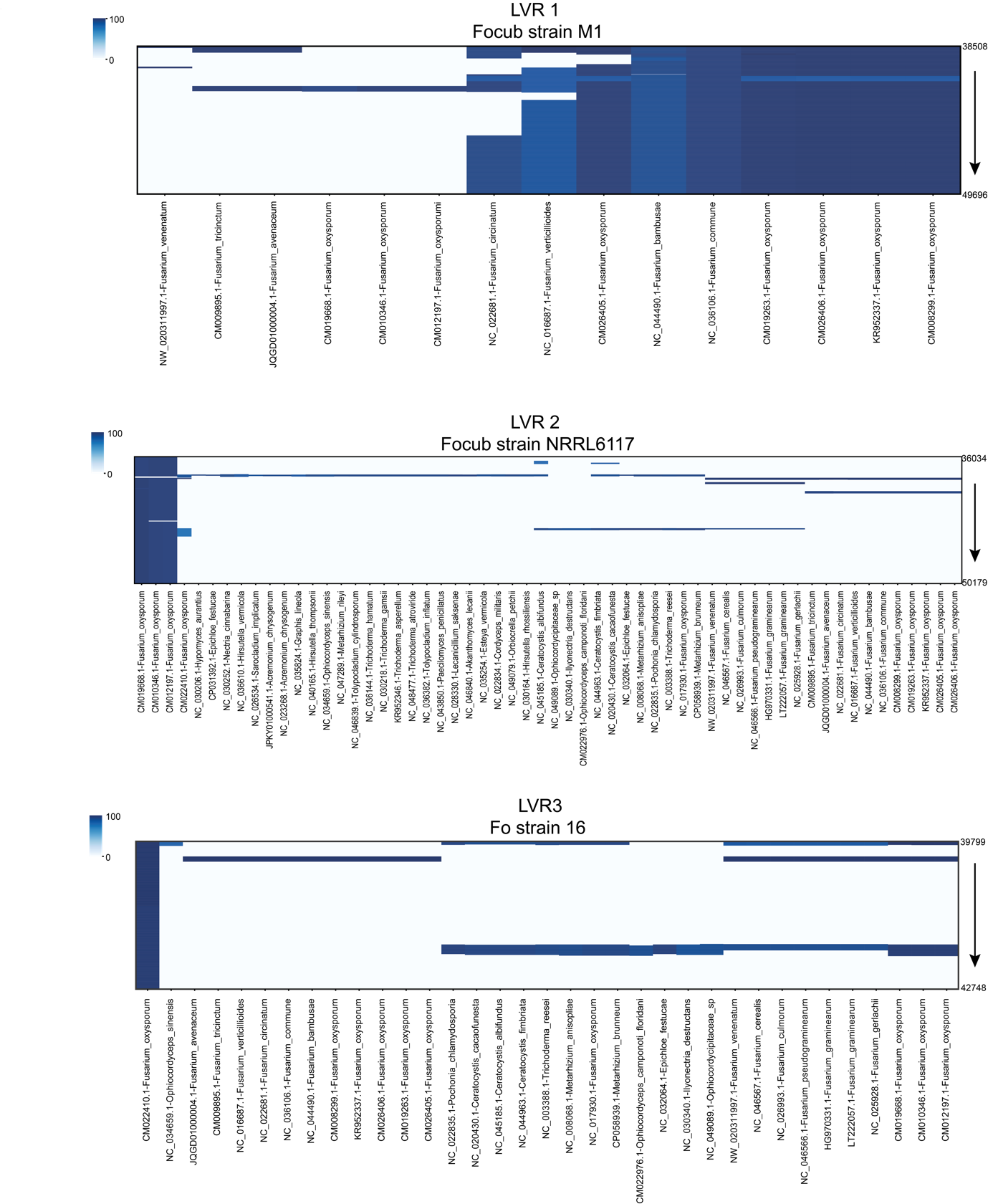

**Figure S4 –** BLAST search of the three long variable regions (**a)** LVR1, (**b)** LVR2, (**c)** LVR3) against 706 fungal mitogenomes (supplementary table 2), indicating that the LVRs are only found in members of the Fusarium genus, outside the Fusarium genus very short hits are found. LVR 1 is found outside of Fusarium oxysporum, whereas LVR 2 and 3 are soley found in members of the *Fusarium oxysporum* species compex. The color indicates the percent identity per genomic region.

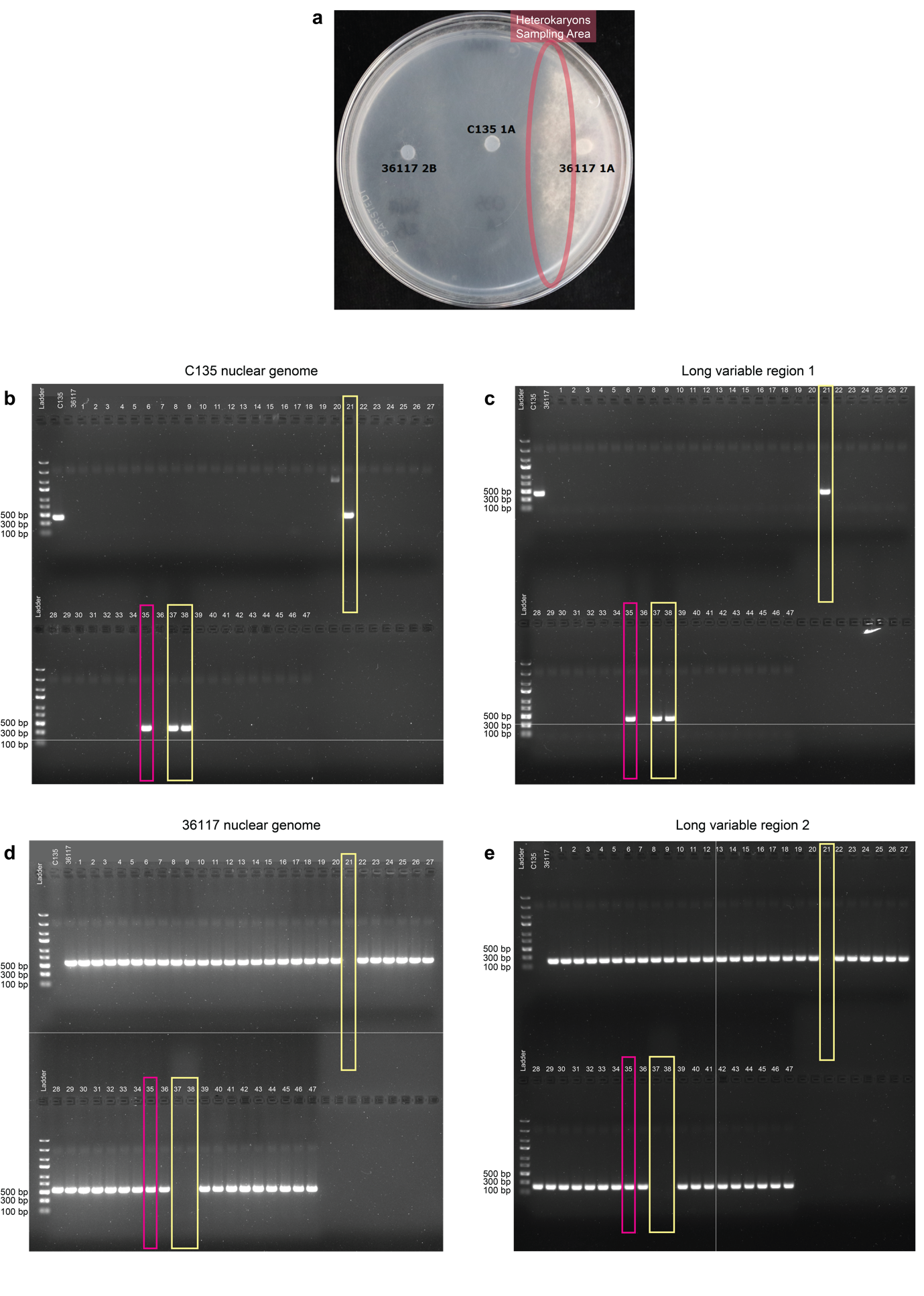

**Figure S5** – PCR identification of the nuclear genome and the mitochondrial long variable region (LVR) upon heterokaryon formation of *Fusarium oxysporum* strain NRRL36117 (LVR 2) and C135 (LVR 1). (**a**) Growth of the C135 and NRRL36117 *nit*-mutants. Upon heterokaryon formation growth of the mutants is restored (red circle). The heterokaryon was sampled and diluted to obtain single spore colonies. 47 colonies have been screen by PCR for **(b)** the C135 nuclear genome primer, **(c)** LVR 1, **(d)** NRRL 36117 nuclear genome and **(e)** the LVR 2 primer. Yellow boxes indicate three spores that are positive for the C135 nuclear genomes and the corresponding LVR1. One out of 47 colonies is still a heterokaryon (purple box) and positive for the NRRL36117 and C135 nuclear genome as well as for both LVRs. No exchange of LVRs is observed upon heterokaryon formation in this experiment.

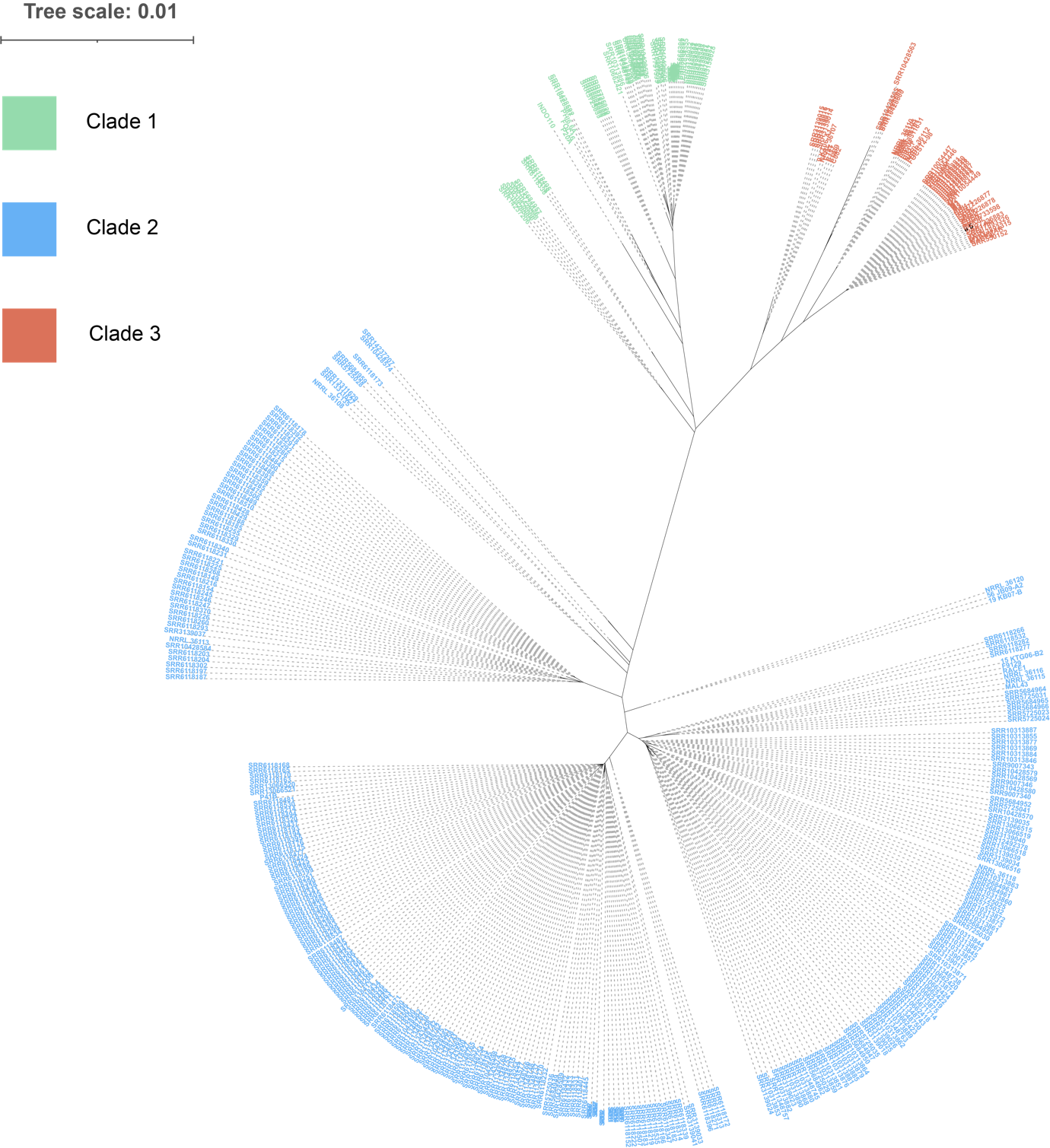

**Figure S6** – Maximum likelihood phylogeny of long variable region 1 (LVR1). Labels are colored according to the nuclear phylogeny. Based on the LVR1 phylogeny, isolates belonging to the same clade group together, suggesting that LVR1 largely follows the nuclear phylogeny.

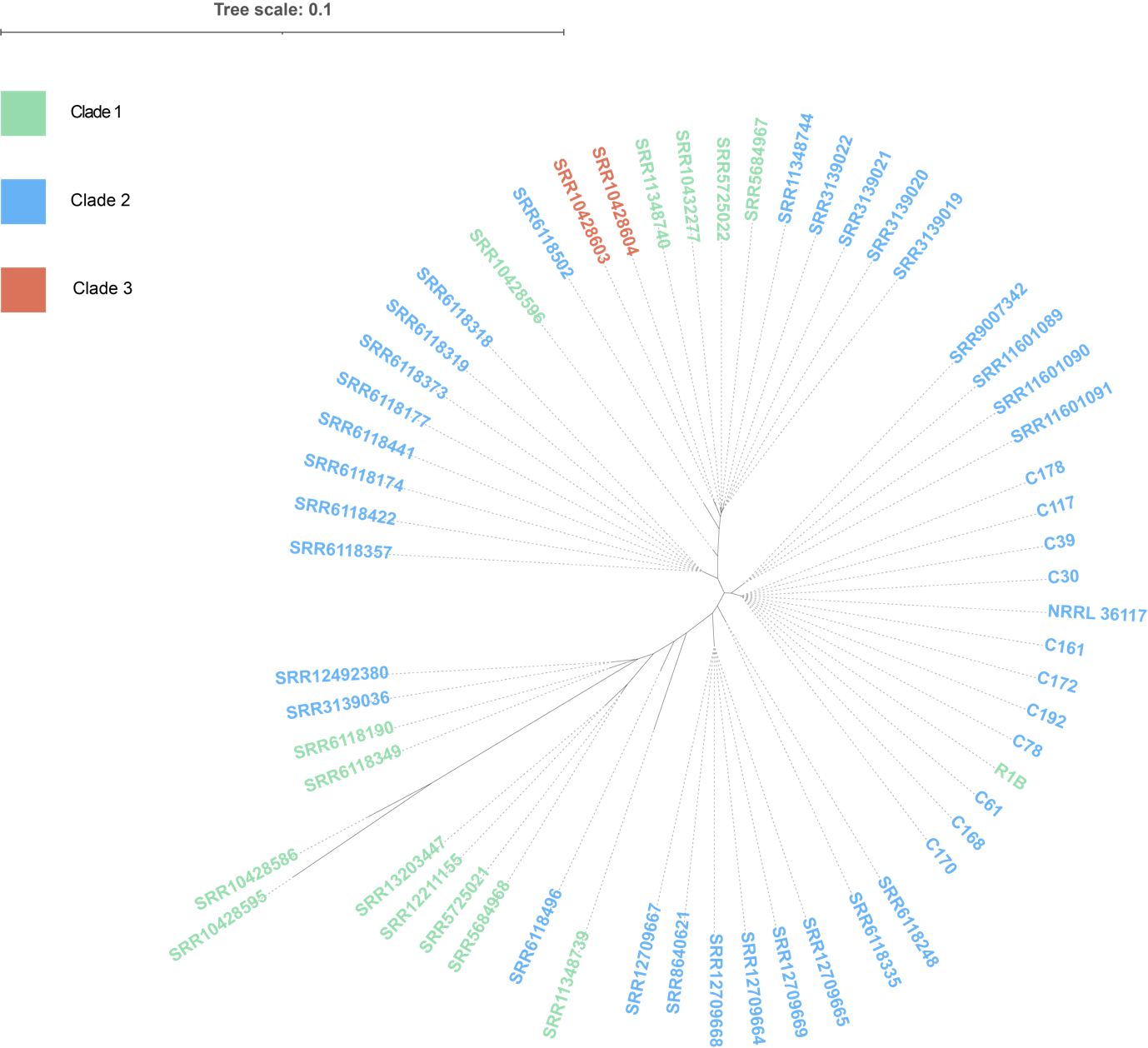

**Figure S7** - Maximum likelihood phylogeny of long variable region 2 (LVR2). Labels are colored according to the nuclear phylogeny. The phylogeny shows that isolates from different clades can share a highly similar LVR2. For example, R1B (Clade 1) groups together with other genomes from Clade 2.

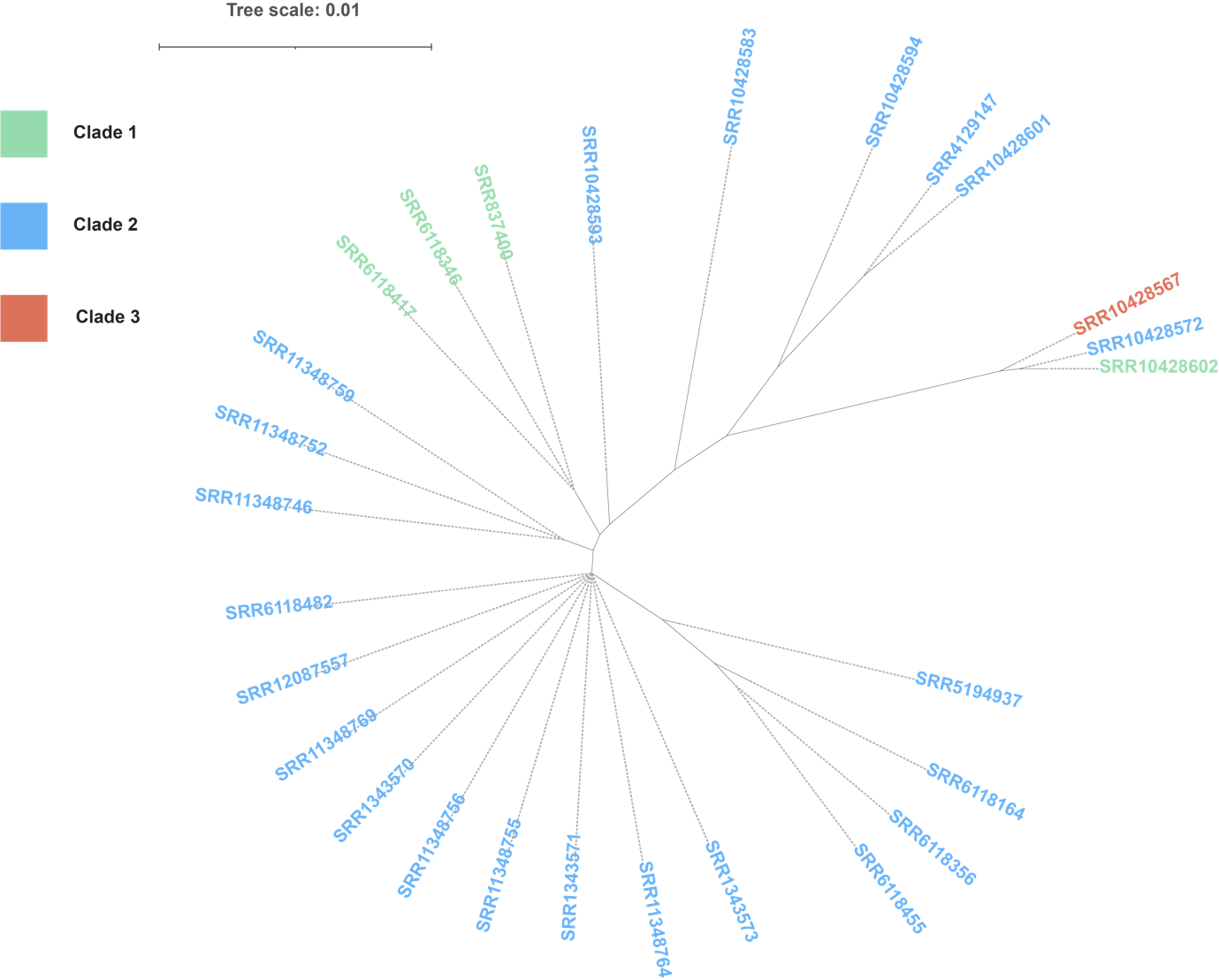

**Figure S8** - Maximum likelihood phylogeny of long variable region 3 (LVR3). Labels are colored according to the nuclear phylogeny. Isolates belonging to a different clade can have a highly similar LVR3. For example, three genomes (SRR10428567; Fo6, SRR10428572; Fo13, SRR10428602; Fo5) of three different clades group together on one branch.
